# Spatiotemporal Dynamics of Molecular Pathology in Amyotrophic Lateral Sclerosis

**DOI:** 10.1101/389270

**Authors:** Silas Maniatis, Tarmo Äijö, Sanja Vickovic, Catherine Braine, Kristy Kang, Annelie Mollbrink, Delphine Fagegaltier, Žaneta Andrusivová, Sami Saarenpää, Gonzalo Saiz-Castro, Miguel Cuevas, Aaron Watters, Joakim Lundeberg, Richard Bonneau, Hemali Phatnani

## Abstract

Paralysis occurring in amyotrophic lateral sclerosis (ALS) results from denervation of skeletal muscle as a consequence of motor neuron degeneration. Interactions between motor neurons and glia contribute to motor neuron loss, but the spatiotemporal ordering of molecular events that drive these processes in intact spinal tissue remains poorly understood. Here, we use spatial transcriptomics to obtain gene expression measurements of mouse spinal cords over the course of disease, as well as of *postmortem* tissue from ALS patients, to characterize the underlying molecular mechanisms in ALS. We identify novel pathway dynamics, regional differences between microglia and astrocyte populations at early time-points, and discern perturbations in several transcriptional pathways shared between murine models of ALS and human *postmortem* spinal cords.

**One Sentence Summary:** Analysis of the ALS spinal cord using Spatial Transcriptomics reveals spatiotemporal dynamics of disease driven gene regulation.

---

Amyotrophic lateral sclerosis (ALS) is a progressive neurodegenerative disease in which symptoms typically manifest at first in distal muscles of a single limb. Symptoms then spread throughout the body, ultimately leading to total paralysis. Loss of locomotor control is the result of denervation of skeletal muscles and motor neuron death. However, the events that initiate disease pathology, and the mechanisms through which pathology spreads remain poorly understood (*1–4*). Mounting evidence indicates that dysfunction in signaling between motor neurons and glia is a key component of the disease (*1, 5, 6*). Inherent limitations of widely available gene expression profiling technologies, such as low throughput or lack of spatial resolution have thus far frustrated efforts to understand how such dysfunction participates in the onset and spread of ALS pathology in the spinal cord. Given the stereotyped cellular organization of the spinal cord and the known importance of intercellular communication in ALS progression, we reasoned that a spatially resolved view of disease driven gene expression changes would be needed to order these events, reveal the relevant sub-populations of cells involved in each stage in disease progression, and characterize the underlying molecular mechanisms that trigger and maintain the disease phenotype.

Spatial Transcriptomics (ST) generates quantitative transcriptome-wide RNA sequencing (RNAseq) data via capture of polyadenylated RNA on arrays of spatially barcoded DNA capture probes (*7–9*). We applied ST to spatially profile gene expression in lumbar spinal cord tissue sections (L3-L5) from SOD1-G93A (ALS) and SOD1-WT (control) mice at pre-symptomatic, onset, symptomatic, and end-stage time points (Table S1). We then applied ST to profile gene expression in *postmortem* lumbar and cervical spinal cord tissue sections from either lumbar or bulbar onset sporadic ALS patients resulting in over 76 thousand spatial gene expression measurements (SGEMs), mapping to ~1200 spinal cord tissue sections of 67 mice, and over 60 thousand SGEMs mapping to 80 *postmortem* spinal cord tissue sections from seven patients (Table S1). An interactive data exploration portal for our ST analysis is available at http://alsst.nygenome.org/.

We annotated each SGEM with an anatomical annotation region (AAR) tag, and used these tags to conduct differential expression analysis and to register the data to a common coordinate system (Figs. S1,S2; Table S1). To estimate gene expression levels accurately and detect significant regional, anatomical, and cell type changes within and between conditions, we formulated a novel hierarchical generative probabilistic model (Data 1). Our model corrects for missing data due to undersampling and bias, which has been a common issue in spatial and single cell RNAseq (scRNAseq) studies. As a result, we reliably quantitated the spatial distribution of 11,138 genes in mouse and 9,624 genes in human spinal cord sections. Furthermore, principal component analysis of the complete mouse SGEM dataset reveals that the majority of the variance is explained by spatial location, disease state, and genotype (Fig. S3), and not by batch effects.

Our analysis recapitulates the specific regional and temporal expression patterns for genes with previously described regional expression profiles (*Mbp, Ebf1*, and *Slc5a7*) (*10–12*) and roles in ALS progression (*Aif1, Gfap*) (*13*). Immunofluorescence (IF) imaging of the protein products of these genes demonstrates spatial concordance with our analysis (Fig. 1; Tables S2,S3). Furthermore, our observations, shown for example by *Fcrls, Aif1, Gfap* and *Aldh1l1*, suggest that microglial dysfunction occurs well before symptom onset, precedes astroglial dysfunction in ALS, and that this early microglial dysfunction is proximal to motor neurons (Table S3; Fig. S4).

**Fig. 1.**
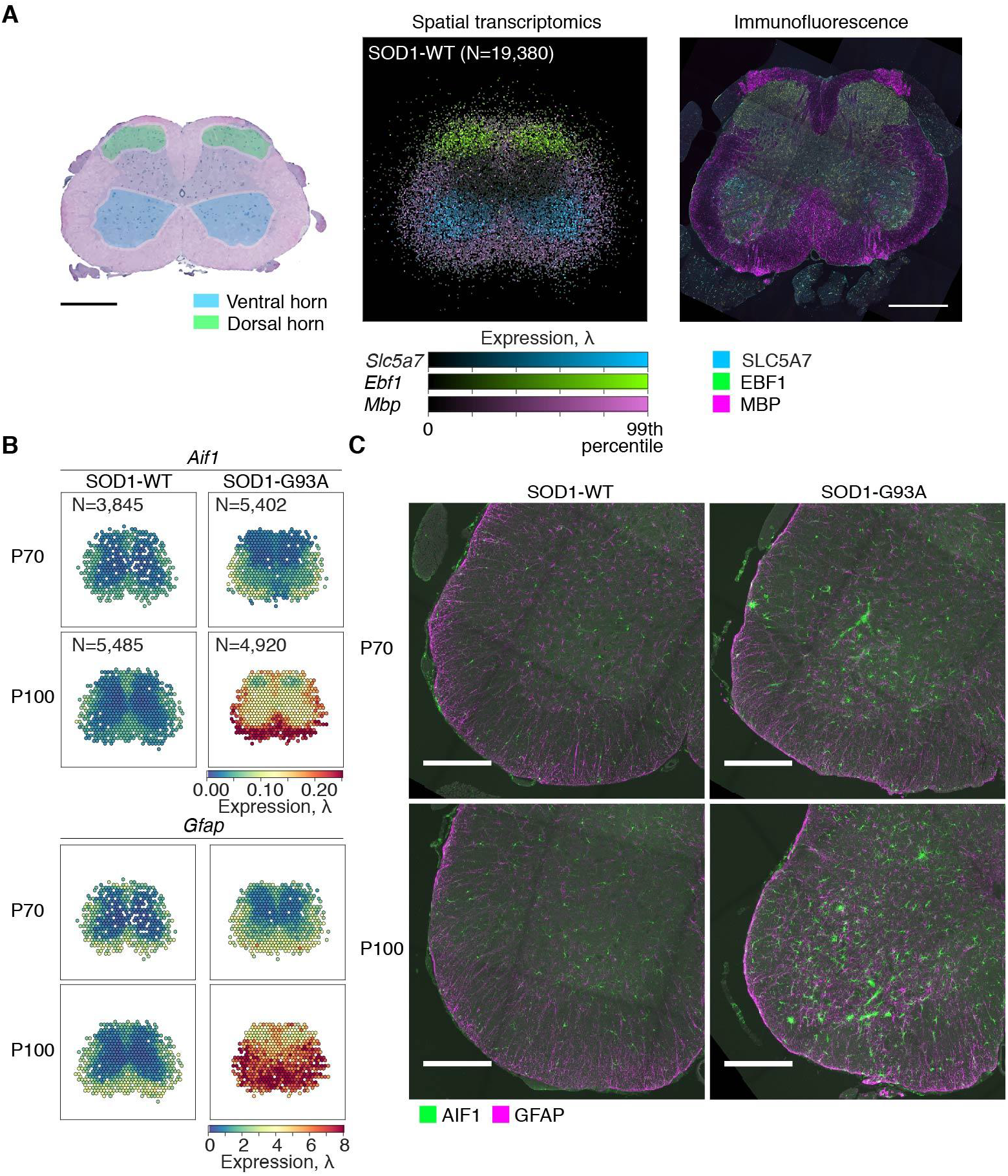
Spatially and temporally resolved gene expression in the mouse spinal cord. (**A**) A schematic diagram of a Hematoxylin and Eosin stained cross-section of mouse lumbar spinal cord with anatomical annotation regions (AARs) overlaid (left panel). Scale bar is 500μm. A multichannel visualization of colocalized spatial mRNA expression (the posterior means of λ) of *Slc5a7* (blue), *Ebf1* (green), and *Mbp* (purple) (middle panel). All analyzed and registered SGEMs (N=19,380) from the control tissue sections were considered. Representative Z maximum projection of 10μm confocal image stack of EBF1 (green), SLC5A7 (blue), and MBP (purple) immunofluorescence in mouse lumbar spinal cord (N=7 animals) (right panel). Scale bar is 500μm. (**B**) Spatial mRNA expression of microglia-expressed *Aif1* and astrocyte-expressed *Gfap* in control and ALS lumbar spinal cords at P70 and P100. (**C**) Representative Z maximum projections of 10*μ*m confocal image stacks of AIF1 (green) and GFAP (magenta) immunofluorescence in control and ALS spinal cords at P70 and P100 (N=12 animals). Scale bars are 250*μ*m.

To further explore the spatiotemporal dynamics of microglial activation, we focused on a mechanism involving TREM2 reported in several neurodegenerative disease models (*6, 14, 15*). TREM2 and TYROBP form a receptor complex that can trigger phagocytosis or modulate cytokine signaling when engaged by membrane lipids, or lipoprotein complexes (*15, 16*). This mechanism also involves *Apoe, Lpl, B2m*, and *Cx3cr1*, and is activated by microglial phagocytosis of apoptotic neurons. *Lpl* is expressed by microglia of the CNS, and aids in the uptake of myelin derived lipids, a process that can modulate microglial activation (*17*). Spatial gene expression analysis suggests a spatiotemporal ordering of this TREM2-mediated mechanism in ALS. We observe that *Tyrobp* is upregulated pre-symptomatically and before *Trem2* in the ventral horn and ventral white matter. Furthermore, *Lpl* and *B2m* are simultaneously upregulated (pre-symptomatically) uniquely in the ventral horn, while *Apoe* and *Cx3cr1* are not (Fig. 2; Table S3; Fig. S4). These latter genes become widely upregulated in spinal cords of symptomatic mice (Table S3). *Apoe* expression is driven by *Trem2* signalling, and is itself a ligand for *Trem2*, and therefore, *Apoe* and *Trem2* expression can act in an autoregulatory loop that can trigger and maintain a phagocytic microglial phenotype (*18*). Collectively, our spatial analysis suggests that TREM2/TYROBP mediated signaling is an early step in disease relevant changes in microglial gene expression. Further, the spatiotemporal ordering of gene expression changes that we observe adds to what can be learned using single cell RNA sequencing of sorted microglia (*6, 14*), demonstrating the value of spatially-resolved high dimensional analyses.

**Fig. 2.**
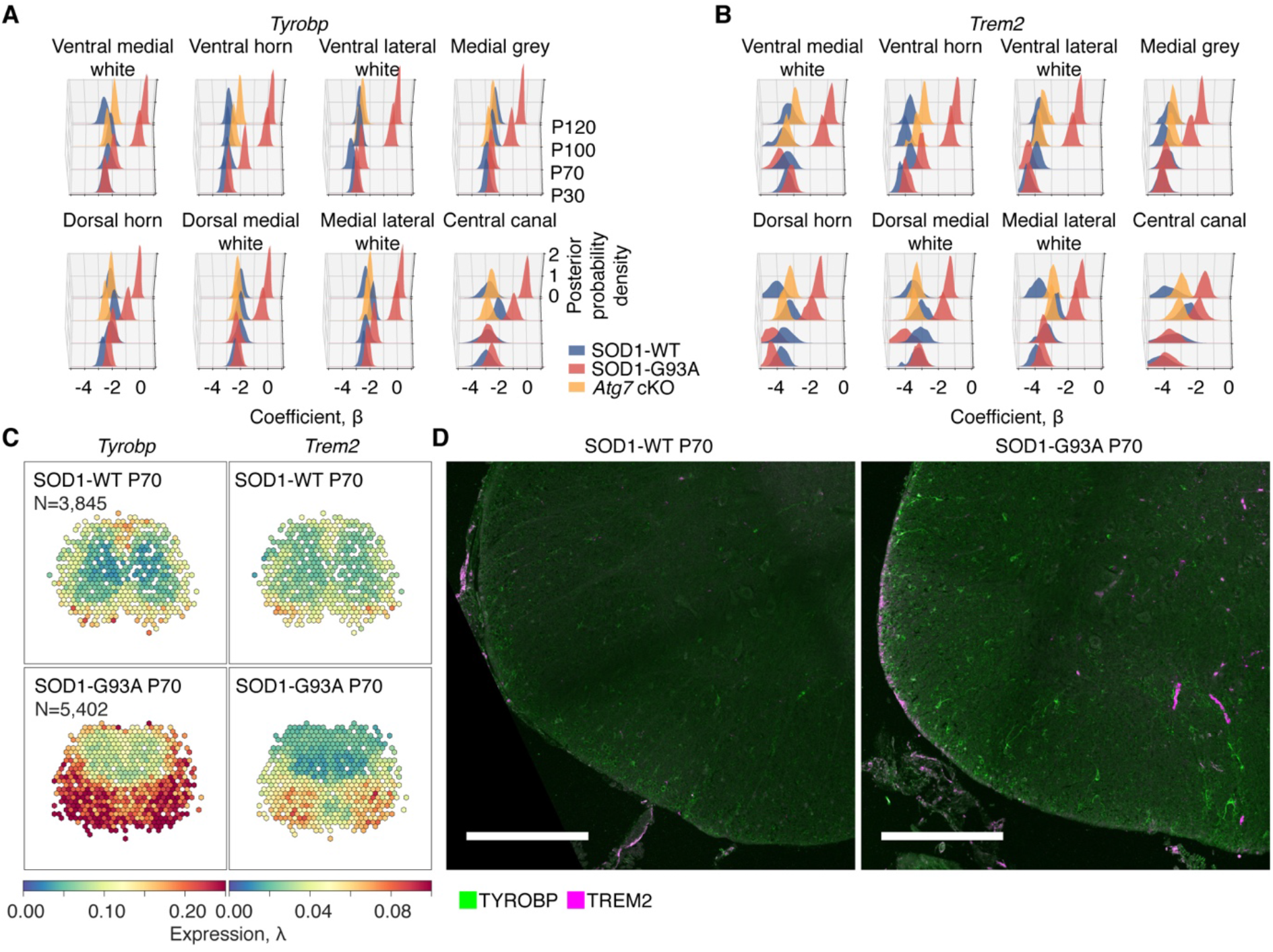
Pre-symptomatic dysregulation of TREM2/TYROBP mediated signaling. (**A**) The posterior distributions of coefficient parameters β of *Tyrobp* of different AARs in control (blue), ALS (red), and *Atg7* cKO (yellow) at P30, P70, P100, and P120. The coefficient parameters β capture offsets of expression (in natural logarithmic space) in distinct AARs across all tissue sections of a given condition. (**B**) As in (**A**), with the focus here on *Trem2*. (**C**) Spatial mRNA expression of *Tyrobp* in control and ALS spinal cords at P70. (**D**) Representative Z maximum projections of 10*μ*m confocal image stacks TYROBP (green) and TREM2 (magenta) immunofluorescence in control and ALS ventral-lateral spinal cords at P70 (N=6 animals). The scale bar is 250*μ*m.

*Trem2* mutations are associated with several neurodegenerative diseases (*16, 19–22*) and, through mTOR signaling in myeloid cells (*16*), *Trem2* expression modulates autophagy. Mutations in several autophagy related genes are associated with ALS (*16*). ST analysis and IF imaging show that genes involved in autophagy and the endolysosomal system are dysregulated in the ALS spinal cord (Fig. S5; Table S3). Ablation of autophagy by conditional knockout of *Atg7* in cholinergic cells, including motor neurons (ChAT-Cre^+/+^; *Atg7*^fl/fl^; SOD1-G93A), leads to earlier symptom onset but prolonged survival in ALS mice (*5*). This manipulation also partially rescues reactive gliosis in ALS mice. To investigate which pathways might link dysfunction in autophagy to gliosis and motor neuron loss in ALS, we applied our methods to these mice (*Atg7* cKO). As expected, we observe that expression of *Gfap* and *Aif1*, and activity of the TREM2 microglial activation axis, are greatly reduced when autophagy is ablated in motor neurons, particularly in AARs distal to motor neuron somata (Table S3).

To better understand disease relevant changes in gene regulation and interactions between cell types, we carried out an unbiased co-expression analysis of our mouse ST data. We identified 31 major co-expression modules (Methods; Fig. 3A; Table S4) of diverse spatiotemporal and pathway activities, a subset of which are affected in Atg7 cKO (Fig. 3B; Figs. S6,S7A; Table S5). Examining these modules in the context of published scRNAseq data (*23*) reveals that many of the modules are comprised of genes preferentially expressed in specific cell types (Fig. S7B). We further grouped the genes of each module based on their cell-type specific expression pattern, resulting in submodules (Methods; Fig. 3C; Tables S6,S7). Submodules that are enriched for a given cell type can display distinct spatiotemporal expression patterns depending on the parent module from which they are derived. Such differences represent functionally distinct subpopulations within that cell type. For example, submodules containing *Prdx6* and *Gfap* (Submodule 8.9) or *Slc7a10* and *Bcan* (Submodule 29.41) represent regional astrocyte subpopulations that behave differently across the course of disease (Fig. S8A). *Slc7a10* has previously been reported as a marker of an astrocyte subpopulation associated with glycinergic signaling in mice (*24*). In ALS animals, the *Prdx6* submodule displays escalating activity as disease progresses, while the *Slc7a10* submodule is attenuated with progression. Intriguingly, the disease related dynamics of the *Prdx6* astrocyte submodule in ALS animals are rescued in *Atg7* cKO, but those of the *Slc7a10* astrocyte submodule are not. Our spatial analysis thus identifies gene expression programs characteristic of regional astrocyte populations (*25*) that display distinct, disease relevant spatiotemporal dynamics and are differentially dependent on cholinergic autophagy.

**Fig. 3.**
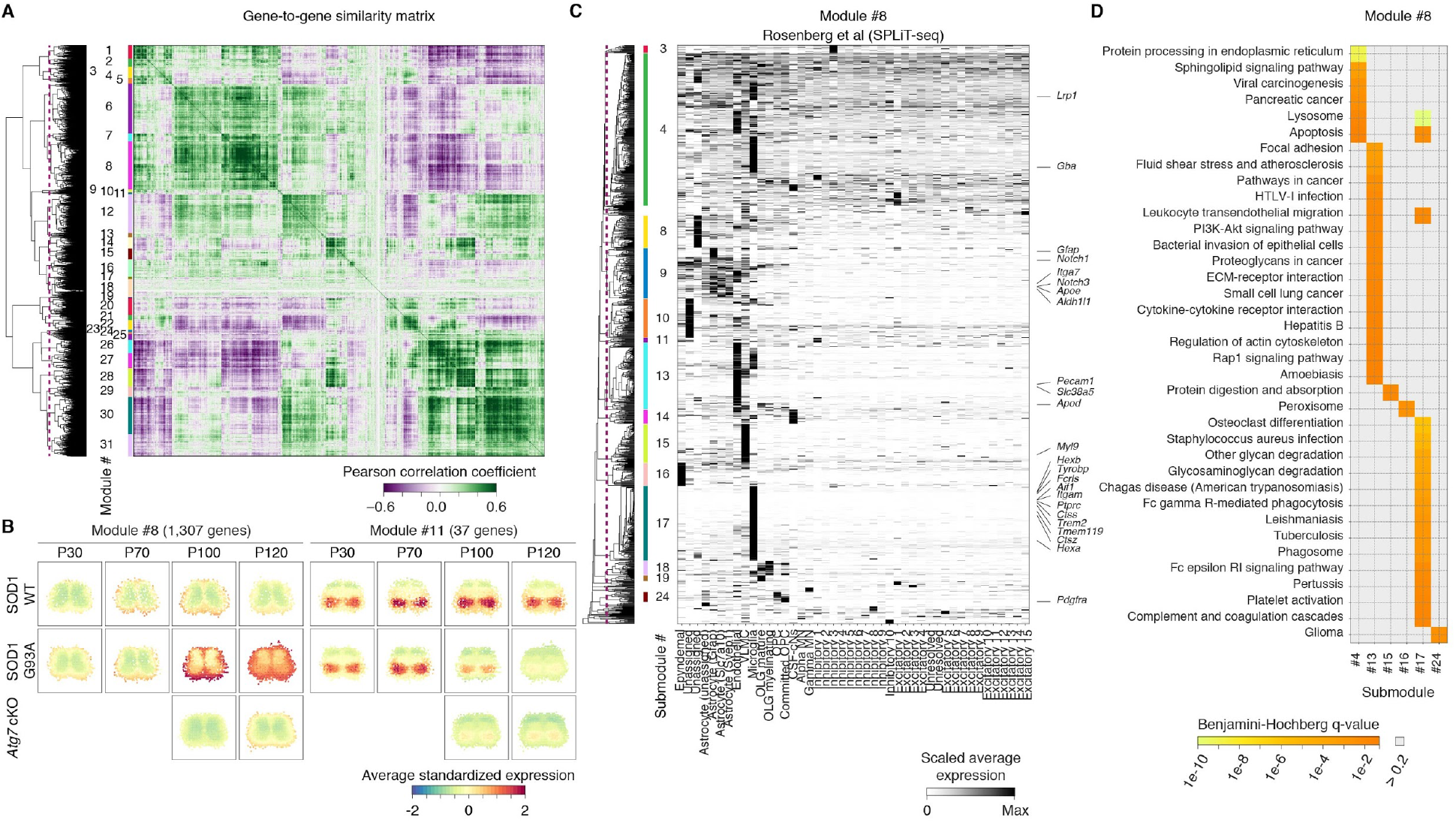
Spatiotemporal dynamics of gene expression during disease progression in ALS. (**A**) Biclustering of the mouse SGEMs reveals spatially and temporally co-expressed genes. The dashed vertical purple line in the dendrogram denotes the break. The identifiers given to the co-expression modules are listed. (**B**) Average spatiotemporal expression dynamics of genes in co-expression modules 8 and 11 are visualized. (**C**) Hierarchical clustering of the genes of co-expression module 8 using independent gene expression data of mouse CNS cell types. The dashed vertical purple line in the dendrogram denotes the break. The identifiers given to the co-expression submodules having at least 10 genes are listed on right of the dendrogram. Selected genes of interest are highlighted on right. (**D**) Analysis of enriched KEGG pathways among the genes of the submodules depicted in (**C**) (one-tailed Fisher’s exact test with Benjamini-Hochberg correction, FDR < 0.1).

This approach also allows us to identify coordinated activities across cell types that reflect major aspects of disease, and to highlight expression programs within distinct cell types associated with these processes (Fig. S7B). In one such example, a microglial expression program containing *Trem2, Tyrobp, Aif1*, and other reactive microglial genes, displays spatiotemporal dynamics reflecting known patterns of progressive gliosis in ALS (Submodule 8.17) (*6*). A second microglial expression program containing *Lrp1* and *Gba* (Submodule 8.4) exhibits spatiotemporal dynamics correlated with those of the *Trem2* microglial submodule 8.17. KEGG analysis of these submodules shows that they are both enriched for lysosomal factors (Fig. 3D). However, while the *Trem2* submodule includes many factors involved in the complement cascade, Fc receptor mediated signaling, and phagocytosis, the *Lrp1* submodule includes many sphingolipid signaling factors. Both the *Trem2* and *Lrp1* submodules exhibit correlated spatiotemporal dynamics with an astrocyte expression program (Submodule 8.9) that includes *Prdx6, Gfap, Aldh1l1, Apoe, Notch1, Notch3* and several TGF-beta superfamily members. *Apoe*, TGF-beta, and Notch signaling all affect astroglial and microglial activities in neurodegeneration, and have been implicated in disease relevant signaling between these cell types (*2, 6, 26, 27*). Lastly, the activity of these microglial and astroglial submodules are correlated with activities of an oligodendrocyte precursor (OPC) expression program including *Pdgfra* (Submodule 8.24) and mature oligodendrocyte submodules (Submodules 8.18 and 8.19) (Fig. S8B). *Pdgfra* expressing OPCs progressively increase in number in the ventral grey matter in ALS mouse spinal cords, and this increase is associated with defects in myelination (*10*). Strikingly, the spatiotemporal pattern of module 8 expression is rescued by ablation of autophagy in cholinergic neurons in *Atg7* cKO mice, demonstrating the key role that neural interactions with module 8 glia play in these activities. Further, this result illustrates the importance of autophagy in ALS pathology. Collectively, the pathway activities encompassed by module 8 reveal signaling within and between cell types during early glial activation in intact tissue, and the mechanisms that maintain and spread the reactive phenotype in the ALS mouse model.

A microglial expression program that includes *Sall1* (Submodule 1.12) is greatly elevated in the white matter of control and presymptomatic ALS animals (Fig. S8C). *Sall1* is expressed by homeostatic microglia, and loss of *Sall1* expression in microglia results in a phagocytic, inflammatory phenotype (*6, 28*). The expression pattern of the *Sall1* submodule illustrates glial interactions that are spatiotemporally and mechanistically distinct from those of module 8. By end stage in ALS animals, this submodule has infiltrated the grey matter, and is attenuated in the white matter. This expression program is coordinated with an astrocyte expression program (Submodule 1.15) that includes *Tlr3*, and an oligodendrocyte program (Submodule 1.8) that includes *Mag, Cnp* and *Mbp* (Fig. S8D). Expression of *Sall1* in this submodule suggests that the late stage expansion of the microglial population responsible for this expression program differs from the clearly reactive microglial populations present in modules 8 and 6 (Fig. S8C). Further, the collective spatiotemporal expression pattern of module 1 in ALS animals is consistent with previously reported late stage defects in myelination (*10*). This view is supported by individual expression dynamics of *Mag, Cnp*, and *Mbp* that we observe. KEGG analysis reveals that genes in module 1 are involved in the Ras signaling pathway, and ether lipid metabolism (Fig. S7A). Notably, *Atg7* cKO does not rescue the dynamics of this expression module, which encompasses the activities of multiple cell types. Thus, spatiotemporal correlation of the *Sall1* microglial, *Tlr3* astroglial, and the oligodendrocyte expression programs in module 1 suggests signaling mechanisms through which glial behavior related to axonal pathology in late stage ALS animals might be coordinated. Taken together, these examples demonstrate the exceptional view of disease related processes that the combination of scRNAseq and ST provides.

Several findings from our mouse study are recapitulated in postmortem spinal cords of sporadic ALS patients. We applied our ST workflow to cervical and lumbar tissue from four patients who presented clinically with bulbar symptom onset, and three patients who presented with lower limb symptom onset. Data from these human samples exhibits expected regional expression patterns, resembling those from the mouse dataset (Fig. S9). Consistent with previous studies (*29*), human spatial data shows variability in gene expression in the anterior horn related to distance from the site of symptom onset (Table S10). For instance, acetylcholinesterase (*ACHE*), the activity of which has been linked to neuromuscular defects in ALS (*16, 19*), shows reduced expression at locations proximal to spinal segments innervating the site of symptom onset (Fig. 4A,B). As with the mouse spatial data, we conducted an unbiased co-expression analysis on the human spatial data, resulting in 28 human expression modules (Fig. S10A,B; Table S8). The spatial mapping patterns for these human expression modules demonstrate AAR characteristic patterns, some of which vary along the rostrocaudal axis (Fig. 4C), or differ between white matter and grey matter (Fig. S10B), or with proximity to site of symptom onset (Fig. S10B). For example, human expression module 3 is attenuated across spinal cord sections at sites proximal to symptom onset (Fig. 4C). Furthermore, this attenuation is most pronounced in the posterior white matter and anterior horns. KEGG analysis shows that human module 3 is enriched for several pathways, including sphingolipid, retrograde endocannabinoid, and WNT signaling (Table S9). This finding, along with several human and mouse submodules that display disease relevant dynamics and that are enriched for sphingolipid signaling pathways, underscores the importance of this mechanism in several aspects of ALS pathology. Indeed, altered glycosphingolipid levels and their metabolism have previously been reported in spinal cords of ALS patients and in murine models of SOD1 ALS (*30, 31*). Modulators of sphingolipid signaling have been proposed as potential therapeutics for ALS, and improve the disease in murine models of ALS (*30, 32*). Our data support this view, detail the dynamics of sphingolipid signaling in multiple cell types, spinal cord regions, and disease stages, and suggest target opportunities for designing therapeutics. Further, the spatiotemporal nature of our data provides insight for the design of treatment strategies for modulating the activity of this pathway based on disease stage. Taken together, this study provides a comprehensive spatiotemporal, transcriptome-wide gene expression dataset with a unique combination of resolution, replication, and biological perturbation. We demonstrate that our procedure scales to real human and clinical settings, and allows us to draw inferences from murine models and test them in clinical samples. As such, we expect the work presented here to be a substantial resource and spur further mapping of the central nervous system and its modes of dysfunction.

**Fig. 4.**
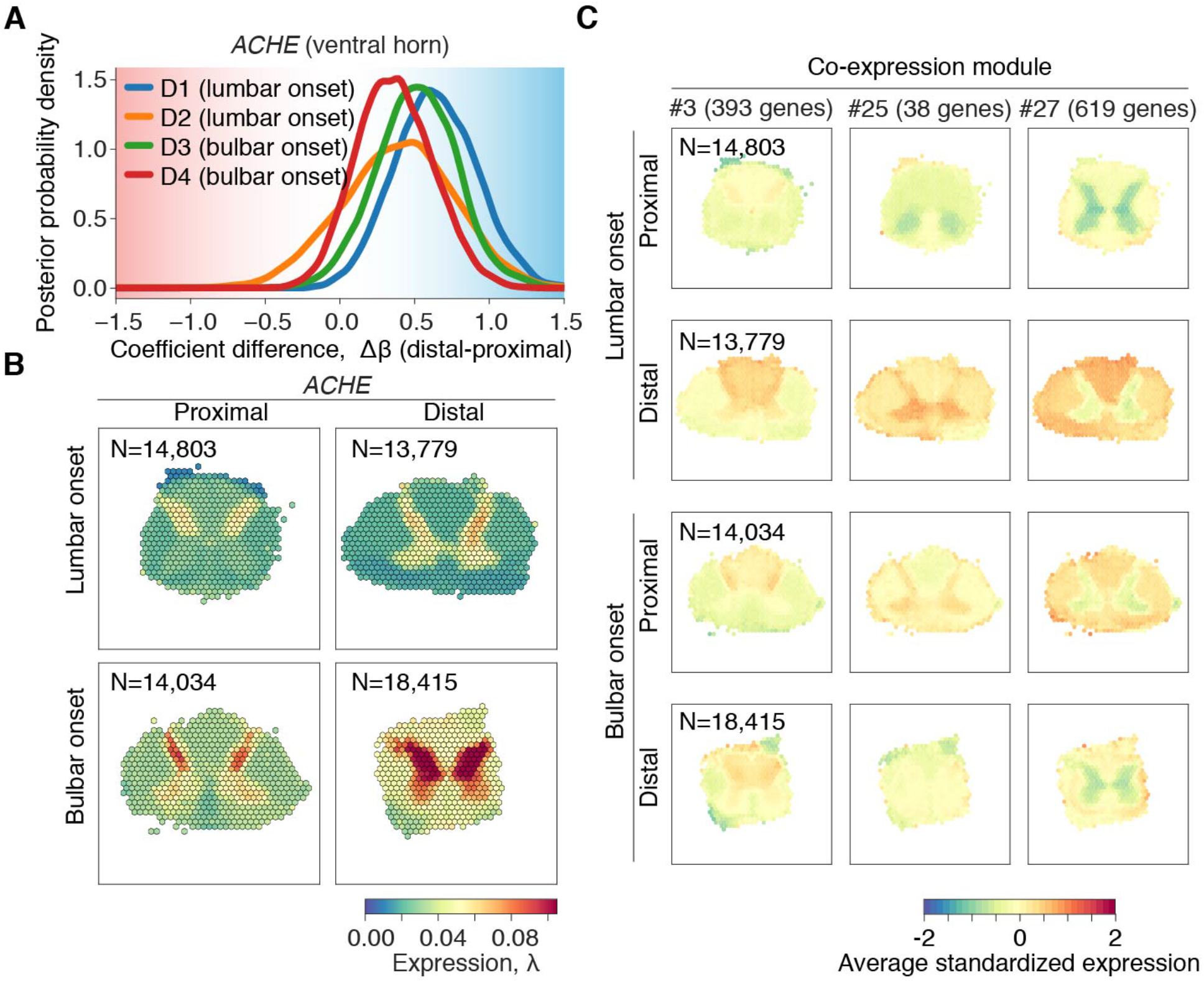
Spatiotemporal transcriptome of human post-mortem spinal cord tissue from ALS patients. (**A**) The posterior difference distributions of the ventral horn coefficients of *ACHE* per patient (D1-D4) are visualized. The differences are calculated between the distal and proximal regions with respect to the onset location. (**B**) Spatial mRNA expression of *ACHE* in human post-mortem lumbar spinal cord and cervical spinal cord tissue sections are visualized. (**C**) Average spatiotemporal expression dynamics of the genes of the human co-expression modules 3, 25, and 27 are visualized.

## Acknowledgments

Human *postmortem* tissue was obtained through the Target ALS Multicenter Postmortem Core. We thank G. Akbalik, J. Gregory, I. Hubbard, D. Kim, and K. Wei for manual anatomical annotation and for manuscript discussions. We thank T. Maniatis for providing *Atg7* cKO mice. We thank Clarapath for the use of their cryostat. We acknowledge the computational resources provided by the Scientific Computing Core of the Flatiron Institute. We thank the NGI Stockholm and SciLifeLab for providing infrastructure support.

## Author contributions

H.P., S.M., and S.V. designed the experiments. S.M. and S.V. performed the experiments, with help from C.B., K.K., M.C., Ž.A.,S.S, and A.M. T.Ä. and R.B. developed and implemented the novel Bayesian generative model and the interactive data exploration portal. S.M., T.Ä., S.V., and D.F. analyzed the data. A.W. implemented the SGEM annotation tool. All authors discussed the results and wrote the manuscript. The project was originally conceived by H.P., with input from all authors throughout experimentation and manuscript preparation.

## Data and materials availability

Raw and processed mouse data and images have been deposited to NCBI’s Gene Expression Omnibus (GEO) Repository under project ID GSE120374. Raw human data have been deposited at New York Genome Center and is available upon request submitted to. A code implementing the used statistical model is available at: https://github.com/tare/Splotch. All processed data and images used in the analyses have been deposited to https://alsst.nygenome.org/.

## Supplementary Materials

Materials and Methods

Tables S1 – S10

Figs S1 – S10

Data 1

References (*33–42*)

